# Trisomy 21 cerebral organoids exhibit Alzheimer’s disease amyloid and apolipoprotein E co-pathologies

**DOI:** 10.64898/2026.07.01.735908

**Authors:** Breanna R. Dooling, Anne Vielle, Esteban M. Lucero, Catherine Rydland, Daphne Quang, Rose Summers, Hector Esquer, Christina Coughlan, Matthew D. Galbraith, Joaquin M. Espinosa, Daniel V. LaBarbera, Heidi J. Chial, Huntington Potter, Aurélie Ledreux, Noah R. Johnson

## Abstract

Adults with Down syndrome (DS) develop Alzheimer’s disease (AD) brain pathology by age 40 due to triplication of the Amyloid Precursor Protein (*APP*) gene on chromosome 21. Inheritance of the apolipoprotein E-e4 (*APOE4)* allele of the *APOE* gene on chromosome 19 remains the greatest genetic risk factor for AD in the typical population, yet its role in DS-associated AD (DS-AD) neuropathogenesis in people with DS is unclear. We generated human induced pluripotent stem cell (hiPSC)-derived neurons, astrocytes, and cerebral organoids (COs) using cells from people with DS and from euploid individuals. Aged DS COs were smaller than aged euploid COs and showed robust amyloid-β neuropathology that was positively correlated with the levels of apoE expression. We then captured extracellular vesicles (EVs) from the conditioned media of COs and observed a decrease in the levels of secreted AD-related proteins, including amyloid, contained within the EVs and in the media from which the EVs were isolated. We also identified distinct neuronal and astrocytic gene expression signatures in DS COs relative to euploid COs, including a set of genes known to interact with both *APOE* and *APP* at the gene and/or protein levels. Lastly, we determined that, despite differences in the expression levels of the specific genes involved, several common pathways were upregulated in T21 hiPSC-derived neurons, astrocytes, and COs, including apoptosis, the endolysosome, and structural stabilization pathways. Taken together, our findings provide novel insights into molecular mechanisms that may contribute to DS-AD and indicate that apoE plays an important role in the disease process.

## Introduction

Alzheimer’s disease (AD) is characterized by extracellular deposits consisting primarily of amyloid-beta (Aβ) peptides and intracellular neurofibrillary tau tangles, which lead to neurodegeneration and cognitive decline [1]. Down syndrome (DS), caused by an extra copy of chromosome 21 (trisomy 21; T21), on which the Amyloid Precursor Protein (*APP*) gene resides, results in additional production of the APP protein and its cleavage products, Aβ peptides, in the brain. Accordingly, all adults with DS develop AD neuropathology by the age of 30-40 years and face a 95% lifetime risk of developing clinical AD [2–4].

Numerous genetic and environmental contributors to AD risk have been identified, yet inheritance of the apolipoprotein E-e4 (*APOE4)* allele of the *APOE* gene on chromosome 19 remains the greatest genetic risk factor for AD [5, 6]. ApoE is the primary cholesterol and lipid transport protein in the central nervous system (CNS) where it is predominantly produced by astrocytes [7]. Three main isoforms of apoE exist in humans—ε2, ε3, and ε4—that differ from each other by a single amino acid at one of two critical residues, causing structural changes in the protein sufficient to affect its function. In the general population, *APOE3* has the highest allelic frequency (∼78%), followed by *APOE4* (∼15%), and *APOE2* (∼7%) [8], with similar allelic frequencies occurring in people with DS [9]. One mechanism by which apoE increases risk for AD is by catalyzing the polymerization of Aβ into fibrils and plaques [10–14], with apoE4 driving earlier and more abundant amyloid plaques. It has been shown that, relative to people who are euploid, apoE levels are increased in the brains of people with DS and are positively correlated with levels of APP and Aβ [15]. These findings suggest that, given the role of apoE as a catalyst for Aβ plaque formation and the increased Aβ load in the DS brain, apoE may play a greater role in the development of DS-associated AD (DS-AD) than in typical AD.

Intercellular communication is paramount to cellular development, neuronal plasticity, and homeostasis in the central nervous system (CNS). Human induced pluripotent stem cell (hiPSC)-derived cerebral organoids (COs), 3D *in vitro* models of the human brain, have been developed to enable studies of the interactions between different CNS cell types [16]. COs include the primary cellular producers of both apoE (i.e., astrocytes) and APP (i.e., neurons) that organize into distinct yet interconnected layers that reflect human brain development. The ability of COs to capture whole-brain activities gives them high physiological relevance compared to other guided brain organoid models [17]. Thus, COs represent valuable models to study the individual and synergistic contributions of neurons and astrocytes to the development of AD phenotypes.

ApoE serves as an important lipid and cholesterol transporter, and current evidence supports an integral role of apoE in extracellular vesicle (EV) biogenesis through endosomal trafficking and membrane-lipid remodeling [18–21]. Increasing evidence shows that EVs are key mediators of the extensive crosstalk between neurons and glia and contribute significantly to the coordinated communication among brain cells [22, 23]. EVs comprise a heterogeneous population of lipid-layer enclosed nanoparticles released by virtually all cell types. They carry proteins, lipids, and nucleic acids that reflect the molecular composition of the cell from which they originate, thereby providing a snapshot of its physiological state [24]. Some EVs, referred to as microvesicles, are formed by outward budding of the plasma membrane while others, often referred to as exosomes, are formed by the inward budding of the endosomal membrane, creating multivesicular bodies that then fuse with the plasma membrane for release into the extracellular space [24]. While exosomes and microvesicles are formed through distinct pathways, they rely on the same machinery, including endosomal sorting complexes required for transport proteins (e.g., ALIX) and tetraspanins [25, 26], making them technically challenging to distinguish.

In neurodegenerative diseases, EVs are increasingly considered to have a dual nature [27]. On one hand, they play a beneficial role by aiding in the clearance of toxic materials in conjunction with autophagic pathways [28]. On the other hand, they can propagate pathology by transferring misfolded or aggregation-prone proteins between cells [29]. Indeed, recent studies have shown that EVs released from neuronal and glial 2D cultures contain Aβ and phosphorylated Tau (pTau), which can be taken up by microglia and astrocytes [30–32]. Additionally, EVs isolated from human plasma can bind to Aβ [33]. Taken together, these results suggest that EVs have the potential to amplify neurodegenerative processes.

Both DS and AD are characterized by abnormalities of the endosomal pathway, including the formation of abnormally numerous and enlarged early endosomes [34, 35]. This disruption is thought to drive aberrant EV secretion, perhaps as a mechanism to clear accumulated toxic materials from the cells [36]. In support of this, emerging studies using human hiPSC-derived brain organoids show that EVs released from these 3D systems carry disease-relevant molecular signatures, including proteins associated with AD [37], supporting their utility as models to investigate EV-mediated mechanisms of pathology.

Here, we sought to determine the roles of apoE3, Aβ, and EVs in the pathogenesis of DS-AD using T21 hiPSC-derived COs. We evaluated the relationship between apoE3 and Aβ protein levels, identified new pathways of differential gene expression, and characterized the EVs secreted from T21 COs to identify their respective contributions to DS-AD phenotypes.

## Results

### T21 COs accumulate increased levels of apoE and Aβ

We generated T21 hiPSC-derived astrocytes, neurons, and COs containing both astrocytes and neurons to establish a complex, physiologically relevant *in vitro* model of the human DS brain. To account for genetic variation, we utilized 15 hiPSC lines derived from 13 individuals with DS or euploid individuals disomic for human chromosome 21 (Hsa21; D21). We confirmed that all hiPSC lines had the expected karyotype (Supplemental Figure 1) and determined that all donors had the *APOE* ε3/ε3 genotype except for one D21 donor (D21-6, Table 1). T21 hiPSC-derived astrocytes were immunopositive for the glial acidic fibrillary protein (GFAP; Figure 1A), as expected, and were of similar size to D21 astrocytes (Figure 1B). T21 neurons were immunopositive for the microtubule-associated protein-2 (MAP2) and for the neuronal nuclei (NeuN) marker (Figure 1C) and were of similar length to D21 neurons (Figure 1D). In COs, GFAP and MAP2 clearly distinguished astrocyte and neuron populations, respectively (Figure 1E). Notably, when developed within COs, T21 astrocytes were significantly smaller than D21 astrocytes (Figure 1F), while T21 neurons were of similar length to D21 neurons (Figure 1G). Finally, all T21 COs, regardless of the age of the donor at time of collection, had a significantly smaller overall volume relative to D21 COs (Figure 1H), as has been reported previously [38].

**Table 1.** Donor information for hiPSC lines.

| ID | Source and external ID | Sex | Cohort | Age (y) | APOE | Notes |
| --- | --- | --- | --- | --- | --- | --- |
| D21-1 | LCI-HTP* Biobank: LC68BC-5 | M | Non-Down syndrome | 39 | ε3/ε3 | Age/sex-matched pair 1 |
| D21-2 | LCI-HTP Biobank: iLC47-4 | M | Non-Down syndrome | 27.5 | ε3/ε3 | Age/sex-matched pair 2;<br>clonal gain of Hsa8 |
| D21-3 | LCI-HTP Biobank: iLC42-3 | M | Non-Down syndrome | 6.1 | ε3/ε3 | Age/sex-matched pair 3 |
| D21-4 | LCI-HTP Biobank: iLD11 #3 | F | Non-Down syndrome | 46.7 | ε3/ε3 | Isogenic pair 1 |
| D21-5 | WiCell: UWWC1-DS2U | M | Non-Down syndrome | 1 | ε3/ε3 | Isogenic pair 2 |
| D21-6 | LCI-HTP Biobank: iC4-4 | F | Non-Down syndrome | 62 | ε2/ε4 | Clonal gain of chromosome 12 |
| D21-7 | LCI-HTP Biobank: iLC62-6 | M | Non-Down syndrome | 27.1 | ε3/ε3 |  |
| D21-8 | LCI-HTP Biobank: iLC67-3 | F | Non-Down syndrome | 28.8 | ε3/ε3 |  |
| T21-1 | LCI-HTP Biobank: LD49BC-3 | M | Down syndrome | 39.1 | ε3/ε3 | Age/sex-matched pair 1 |
| T21-2 | LCI-HTP Biobank: iLD23-2 | M | Down syndrome | 23.2 | ε3/ε3 | Age/sex-matched pair 2 |
| T21-3 | LCI-HTP Biobank: iLD19-4 | M | Down syndrome | 6.2 | ε3/ε3 | Age/sex-matched pair 3 |
| T21-4 | LCI-HTP Biobank: iLD11(2)-1 | F | Down syndrome | 46.7 | ε3/ε3 | Isogenic pair 1 |
| T21-5 | WiCell: UWWC1-DS1 | M | Down syndrome | 1 | ε3/ε3 | Isogenic pair 2 |
| T21-6 | LCI-HTP Biobank: iLD7-6 | M | Down syndrome | 30.3 | ε3/ε3 |  |
| T21-7 | LCI-HTP Biobank: iLD2-3 | F | Down syndrome | 25.1 | ε3/ε3 |  |

\* Linda Crnic Institute Human Trisome Project

**Figure 1.**
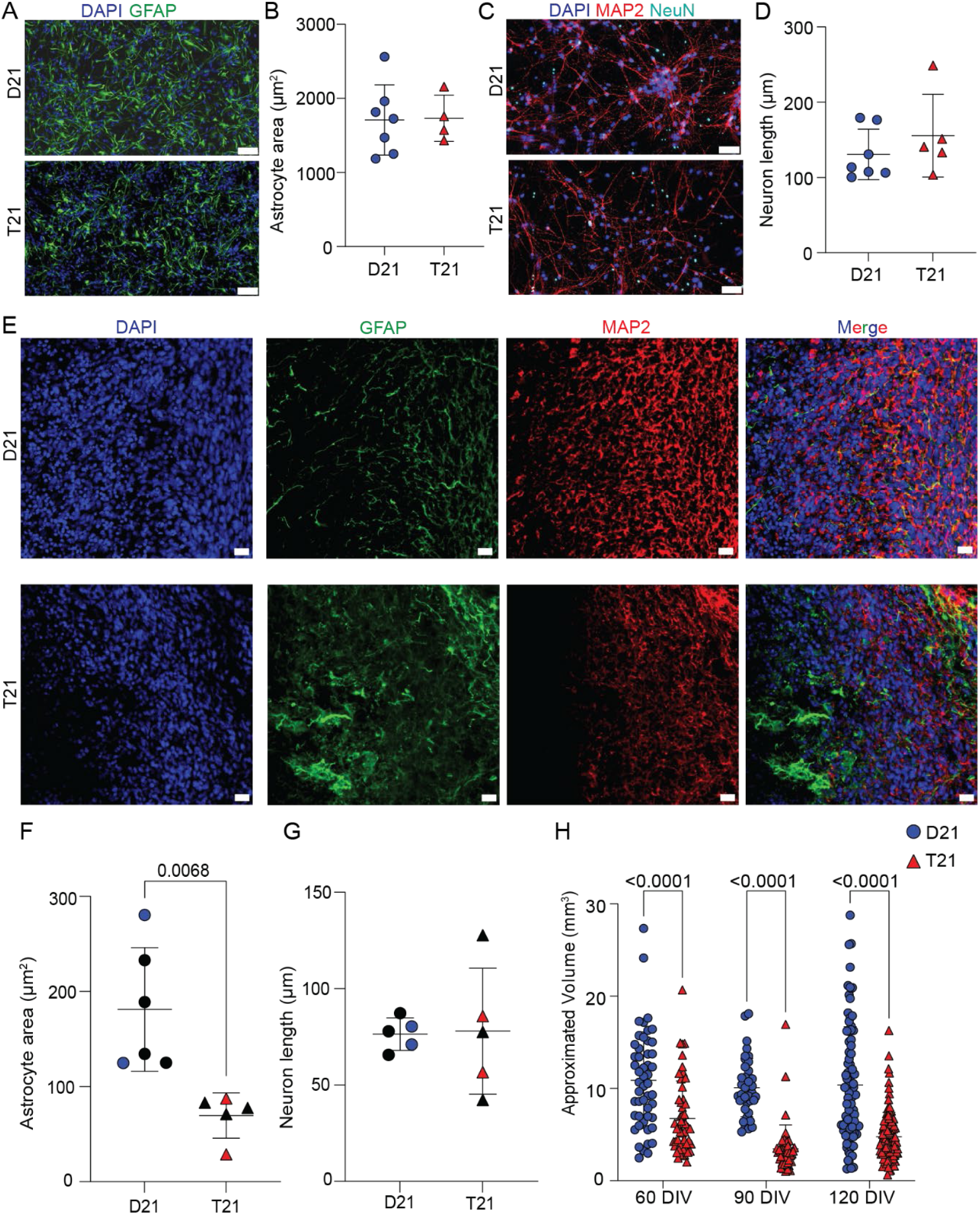
Development within a CO impacts T21 astrocyte and neuron morphology. (A) Representative images of GFAP and DAPI staining of D21 and T21 hiPSC-derived astrocytes. Scale bars are equal to 100 μm. (B) D21 and T21 hiPSC-derived astrocyte area measured using QuPath. Each point represents the average area of 3–6 cells of a unique cell line, with the represented cell lines being D21-1, D21-2, D21-3, D21-4, D21-6, D21-7, D21-8, T21-3, T21-5, T21-6, and T21-7. (C) Representative images of MAP2, NeuN, and DAPI staining of D21 and T21 hiPSC-derived forebrain neurons. Scale bars are equal to 100 μm. (D) D21 and T21 hiPSC-derived forebrain neuron length (as measured from end to farthest end) quantified using QuPath. Each point represents the average length of 3–6 cells of a unique cell line, with the represented cell lines being D21-1, D21-2, D21-3, D21-4, D21-6, D21-7, D21-8, T21-3, T21-4, T21-5, T21-6, and T21-7. (E) DAPI, GFAP, and MAP2 staining in aged D21 and T21 COs with days *in vitro* (DIV) = 90–120. Scale bars are equal to 100 μm. (F) Area of astrocytes measured using GFAP staining within COs quantified in QuPath. Each point represents the average for an individual CO, with two different starting hiPSC lines per genotype represented by either solid black (D21-5 and T21-5) or blue/red colored (D21-8 and T21-1) shapes. (G) Neuron length within COs measured using MAP2 staining within COs quantified in QuPath. Each point represents the average for an individual CO, with two different starting hiPSC lines per genotype represented by either solid black (D21-5 and T21-5) or blue/red colored (D21-8 and T21-1) shapes. (H) Approximated CO volume at three different timepoints in aging measured by the acquisition of 4X light microscopy images and diameter tracing in ImageJ. Each point represents an individual CO. The ages of the COs (ranging from 60–120 DIV) and their chromosome 21 copy number (D21 or T21) were each found to be independent, significant sources of variation, with *P* = 0.004 and *P* < 0.0001, respectively. All results are presented as mean ± standard deviation.

The T21 COs developed substantial Aβ neuropathology, observable as extracellular puncta and large-scale aggregates, whereas the D21 COs developed only minimal Aβ puncta that appeared to be mostly intracellular (Figure 2A). Immunohistochemical (IHC) quantification confirmed that T21 COs had much greater Aβ neuropathology than did D21 COs (Figure 2B). These findings align with previous studies showing that triplication of *APP* in T21 COs results in increased production of Aβ peptides relative to D21 COs [39, 40].

**Figure 2.**
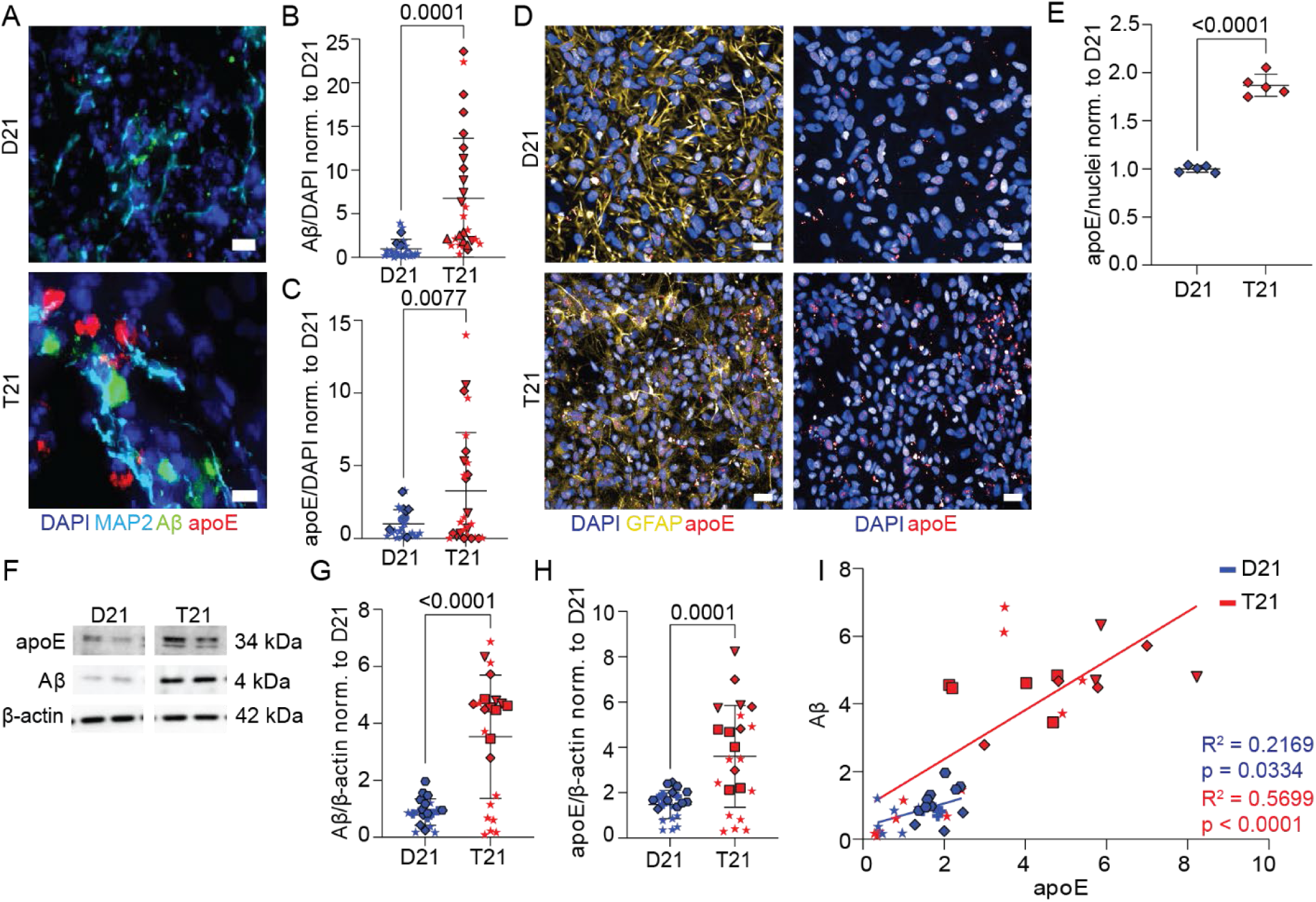
Aβ and apoE protein levels are correlated in aged T21 COs. (A) Representative IHC images of DAPI, MAP2, Aβ, and apoE in aged (DIV = 90–120) D21 and T21 COs with scale bars equal to 10 μm. (B) Quantification of total Aβ IHC staining (normalized to total DAPI staining and D21 mean) in aged D21 (n = 3) and T21 (n = 3) COs. (C) Quantification of total apoE IHC staining (normalized to total DAPI staining and D21 mean) in aged D21 and T21 COs. (D) Representative IHC staining of DAPI, GFAP, and apoE in D21 and T21 hiPSC-derived astrocytes with scale bars equal to 50 μm. (E) Quantification of total IHC staining of apoE (normalized to nuclei count and D21 mean) in D21 (n = 1) and T21 (n = 1) hiPSC-derived astrocytes. (F) Representative images of immunoblot bands for apoE, Aβ monomers, and β-actin. (G) Quantification of monomeric and oligomeric Aβ immunoblot banding (4 kDa – 56 kDa; normalized to sample-specific 42 kDa β-actin bands and D21 mean) in aged D21 (n = 4) and T21 (n = 4) COs. (H) Quantification of 34 kDa apoE immunoblot banding (normalized to sample-specific 42 kDa β-actin bands and D21 mean) in aged D21 (n = 4) and T21 (n = 4) COs. (I) Scatter plot with linear regression for the relationship between Aβ and apoE as each quantified in D21 and T21 immunoblots. The cell lines used in these experiments are D21-1 and T21-1 (diamonds), D21-4 (hexagons), D21-5 and T21-5 (stars), D21-6 (plus signs), D21-8 (circles), T21-6 (upside down triangles), and T21-7 (squares). All results are presented as mean ± standard deviation.

As it does in typical age-associated AD, apoE co-accumulates within amyloid deposits in the brains of individuals with DS-AD [15]. Whether such co-aggregation occurs in T21 COs has not been investigated. In our experiments, we observed large aggregates of apoE in T21 COs but not in D21 COs (Figure 2A), which was confirmed by IHC quantification (Figure 2C). Consistent with these observations, we also observed greater apoE accumulation in T21 hiPSC-derived astrocytes relative to D21 hiPSC-derived astrocytes (Figure 2D), which was confirmed by IHC quantification (Figure 2E). Notably, while there was some spatial overlap between Aβ and apoE signals, most apoE aggregates in T21 COs were near, but not directly colocalized with, Aβ aggregates (Figure 2A). These data suggest that although common mechanisms may be responsible for the accumulation of both proteins, apoE is not passively deposited within amyloid plaques in T21 COs.

To confirm our observations, we performed immunoblot analyses using CO tissue lysates. We observed apoE and Aβ monomer banding at the expected molecular weights (Figure 2F). Consistent with the IHC analyses, T21 COs contained significantly higher levels of Aβ (Figure 2G) and apoE (Figure 2H) relative to D21 COs. Furthermore, Aβ and apoE levels were positively correlated with each other in both D21 COs (R^2^ = 0.2169, *P* = 0.0334) and T21 COs (R^2^ = 0.5699, *P* < 0.0001; Figure 2I). Importantly, Aβ and apoE levels appeared to be consistently correlated in COs derived from all four T21 hiPSC lines tested, suggesting that this phenomenon is not influenced by the genetic background of the hiPSC donor.

### T21 COs exhibit dysregulated EV secretion

The observed alterations in apoE protein expression suggest disturbances in intercellular expression and lipid homeostasis in T21 COs. Therefore, we next sought to determine whether these changes extend to the extracellular compartment by characterizing EVs released by the COs. EVs were isolated from CO conditioned media and characterized using nanoparticle tracking analysis (NTA) on a ZetaView instrument, transmission electron microscopy (TEM), and immunoblotting (Figure 3A). Membrane-bound vesicles exhibiting the characteristic cup-shaped morphology were visualized by negative stain TEM (Supplemental Figure 2A). Fluorescent NTA demonstrated that tetraspanin-positive EVs derived from both T21 and D21 COs were predominantly within the expected size range of 50–150 nm (Supplemental Figure 2B). EV-associated markers including ALIX, CD63, and CD9/CD81 were present in both T21 and D21 CO-derived EVs and were validated by immunoblotting (Supplemental Figure 2C). There was a trend towards a decreased in EVs secreted per CO by T21 COs (Supplemental Figure 2D).

**Figure 3.**
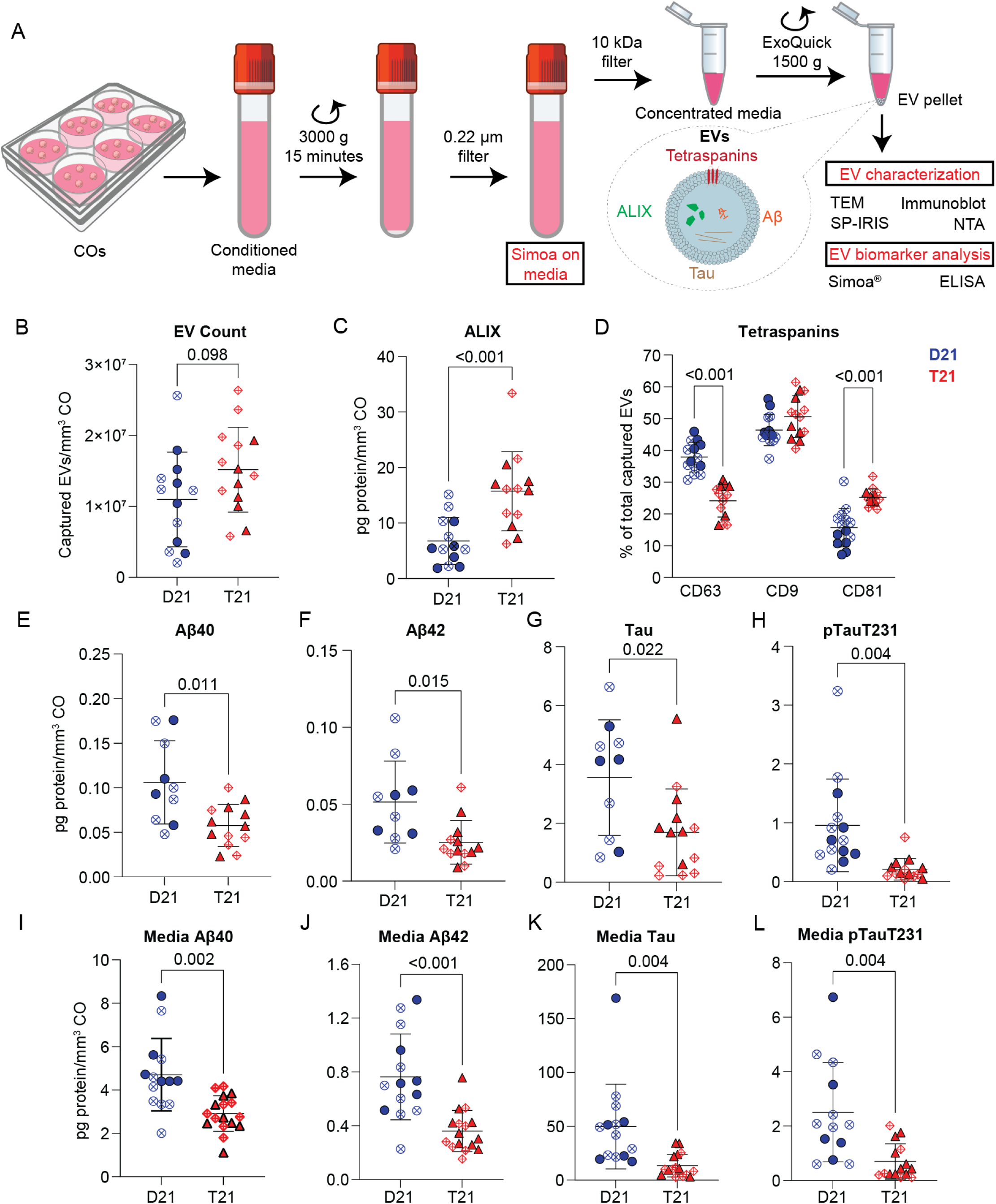
EVs and their cargo are altered in T21 COs compared to D21 COs. (A) Schematic overview of the experimental process used to analyze EV structure composition and associated biomarkers. EVs were isolated from CO conditioned media and EV structure and composition were analyzed using transmission electron microscopy (TEM), immunoblotting to detect EV-associated markers (i.e., ALIX, CD63, and CD9/CD81), single-particle interferometric reflectance imaging sensor-based (SP-IRIS) analyzer (ExoView R200, NanoView Biosciences) to examine tetraspanin distribution, and nanoparticle tracking analysis (NTA) on a ZetaView instrument to determine EV size and number. EV-associated biomarkers were measured using Simoa^®^ technology (SR-X benchtop analyzer, Quanterix) and ELISA. (B) Number of tetraspanin-positive EVs released by D21 and T21 COs normalized to one unit volume of CO. (C) Levels of ALIX measured by ELISA and normalized to one unit volume of CO (mm^3^) by using the median volume of one CO and the numbers of COs for each group. (D) Relative percentages of EVs captured on each tetraspanin-coated chip using the SP-IRIS analyzer. Our results indicate that EVs released by T21 COs contain a significantly lower proportion of CD63-positive EVs and a significantly higher proportion of CD81-positive EVs. Levels of AD-associated proteins, including (E) Aβ40, (F) Aβ42, (G) total Tau, and (H) pTauT231, in the CO-derived EVs were measured using Simoa^®^ technology on the SR-X benchtop analyzer (Quanterix) and normalized to one unit volume of CO (mm^3^). Levels of (I) Aβ40, (J) Aβ42, (K) total Tau, and (L) pTau T231 in CO conditioned media after EV isolation were measured on the SR-X benchtop analyzer (Quanterix) and normalized to one unit volume of CO. All results are presented as mean ± standard deviation.

EVs isolated from CO conditioned media were next characterized by surface tetraspanin profiling on the ExoView instrument. Because T21 COs were significantly smaller than their D21 counterparts, EV measurements were normalized to the number of captured EVs to one unit volume (mm^3^) of COs by considering both the number of COs from which the media was collected and the median CO volume for each age and genotype (Figure 1K). We observed the same patterns across our results with normalization to 10^6^ EV (Supplemental Figure 2E–F). There was a trend towards an increased normalized number of EVs in T21 versus D21 CO conditioned media, but these numbers were not statistically significantly different (Figure 3B). However, the levels of ALIX, a luminal EV marker associated with EV biogenesis, were significantly increased in T21 EVs relative to D21 EVs (Figure 3C). Furthermore, T21 EVs displayed an altered tetraspanin profile with a lower proportion of CD63^+^ EVs and a higher proportion of CD81^+^ EVs (Figure 3D). Taken together, our findings suggest that T21 COs exhibit alterations in EV biogenesis and/or endosomal trafficking pathways.

### T21 CO-derived EVs contain reduced levels of Aβ and pTau

To determine whether EVs released from T21 COs contain altered levels of AD-associated cargo proteins, we used Simoa^®^ technology on the SR-X benchtop analyzer (Quanterix) to measure the levels of Aβ40, Aβ42, total Tau, and pTauT231 in lysed EVs. We found that EVs released from T21 COs contained significantly lower levels of AD-associated proteins relative to EVs released from D21 COs. Specifically, the levels of Aβ40, Aβ42, and total Tau were approximately two-fold lower in T21 EVs than in D21 EVs (Figure 3E–G). Notably, the levels of pTauT231 in T21 EVs were approximately 5-fold lower than in D21 EVs (Figure 3H), which corresponded to a substantially greater reduction than the other measured proteins.

The levels of the same AD-associated proteins were also measured in the conditioned media from which the EVs were isolated. Similar to our lysed EV-associated measurements, we found that the levels of Aβ40, Aβ42, total Tau, and pTauT231 were significantly lower in conditioned media from T21 COs after EV isolation compared to media derived from D21 COs (Figure 3I–L). Furthermore, we found that the Aβ42/Aβ40 ratio was significantly lower in T21 CO conditioned media after EV isolation compared to media derived from D21 COs (Supplemental Figure 2G). Taken together, these findings suggest that reduced EV-mediated clearance of pathogenic proteins may be one possible explanation for the accumulation of AD-associated proteins within T21 COs.

### T21 COs contain a distinct transcriptomic signature of disease

To interrogate potential mechanisms underlying Aβ and apoE neuropathologies caused by T21, we performed bulk RNA sequencing of neurons, astrocytes, and COs. Using DESeq2, we identified a total of 481 differentially expressed genes (DEGs; 272 upregulated and 209 downregulated) across our three T21 sample types relative to D21 samples (Figure 4A–C). Of the upregulated genes, 100 were located on Hsa21 (representing 44.4% of all Hsa21 genes), and no Hsa21 genes were downregulated. Notably, only 36 T21 DEGs (7.7% of all T21 DEGs) were present in two or more sample types (Figure 4D), while the vast majority of DEGs were only found in one sample type (i.e., neurons, astrocytes, or COs). Indeed, only 11 DEGs in T21 COs (10.5% of all DEGs in T21 COs) were also present in T21 neurons or astrocytes. Eight DEGs (five upregulated and three downregulated) were present in both T21 COs and neurons and only one DEG, *H2BC5*, was upregulated in both T21 COs and astrocytes. Only two DEGs were conserved across all three T21 sample types: *PIGP* and *PFKL*, both of which were upregulated. All other DEGs were unique to a single sample type (95 in T21 COs, 296 in T21 neurons, and 54 in T21 astrocytes; Figure 4D). Among the most upregulated Hsa21 genes in T21 COs, we observed minimal expression changes in T21 neurons and astrocytes (Figure 4E). Likewise, the most upregulated and downregulated non-Hsa21 genes in T21 COs were largely the same in T21 neurons and astrocytes (Figure 4F). Lastly, we observed no difference in *APOE* mRNA expression levels between T21 and D21 in COs or astrocytes (Supplemental Figure 3A–B). Taken together, the unique signatures of neurons and astrocytes in comparison to those of COs suggest that neuron-astrocyte interactions are likely to play key roles in the brain’s response to T21, which was recapitulated in our CO model system.

**Figure 4.**
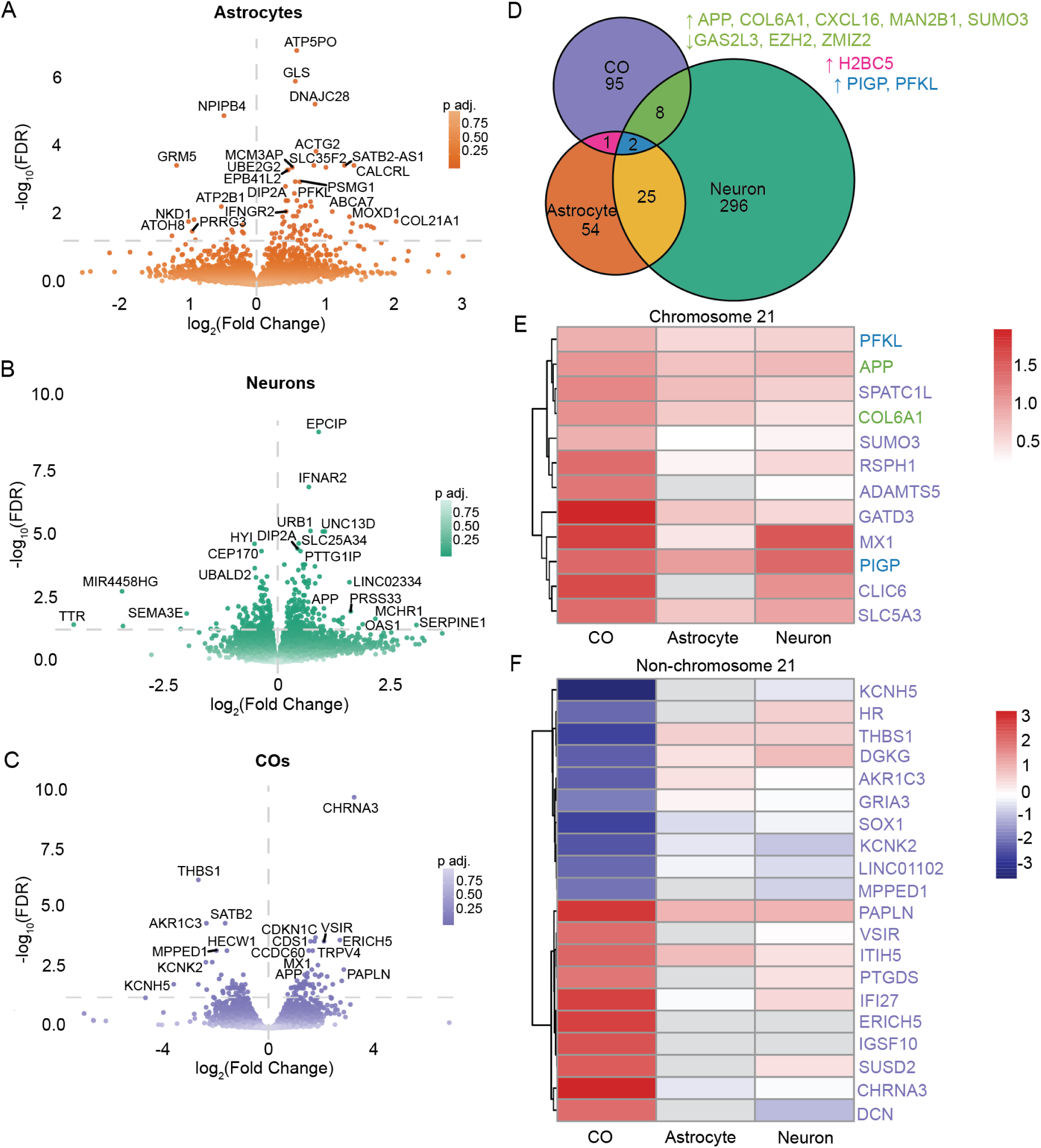
T21 results in unique transcriptomic differences between COs and hiPSC-derived astrocytes and neurons. (A) Volcano plot of differentially expressed genes (DEGs) identified by DESeq2 between D21 and T21 hiPSC-derived astrocytes. (B) Volcano plot of DEGs identified by DESeq2 between D21 and T21 hiPSC-derived forebrain neurons. (C) Volcano plot of DEGs identified by DESeq2 between D21 and T21 aged (DIV = 90–120) COs. (D) Venn diagram showing total counts of DEGs in T21 COs, hiPSC-derived astrocytes, and hiPSC-derived forebrain neurons. Genes that were significantly different in more than one sample type are counted in overlapping regions and listed in the corresponding color. (E) Heat map of log2 fold changes per sample type for the genes on chromosome 21 with the top 12 greatest log2 fold change values in the CO DEG list. Label color indicates Venn diagram bin. (F) Heat map of log2 fold changes per sample type for the genes not on chromosome 21 with the top 12 greatest log2 fold change values in the CO DEG list. Label color indicates Venn diagram bin. The cell lines used in these experiments are D21-1, D21-2, D21-3, D21-4, D21-6, D21-7, D21-8, T21-3, T21-4, T21-5, T21-6, and T21-7.

### T21 COs exhibit dysregulation of AD-associated gene expression pathways

We next sought to identify cell type-specific contributions to gene expression patterns present in T21 COs. First, DEGs were mapped to biological domains using an AD-specific gene ontology resource [41]. In all three T21 sample types (i.e., neurons, astrocytes, and COs), most upregulated DEGs were associated with proteostasis (Figure 5A–C), and outsized proportions were also associated with apoptosis, endolysosomal function, and structural stabilization (Figure 5D). In T21 COs, a large proportion of upregulated genes were associated with tau homeostasis and oxidative stress, pathways that were also highly expressed in T21 neurons (Figure 5D). Among other pathways increased in T21 COs, lipid metabolism was primarily upregulated in T21 astrocytes rather than in neurons (Figure 5D). These pathways may reflect the neuronal and astrocytic contributions or responses to T21-associated pathologies observed in COs including increased Aβ and apoE protein levels.

**Figure 5.**
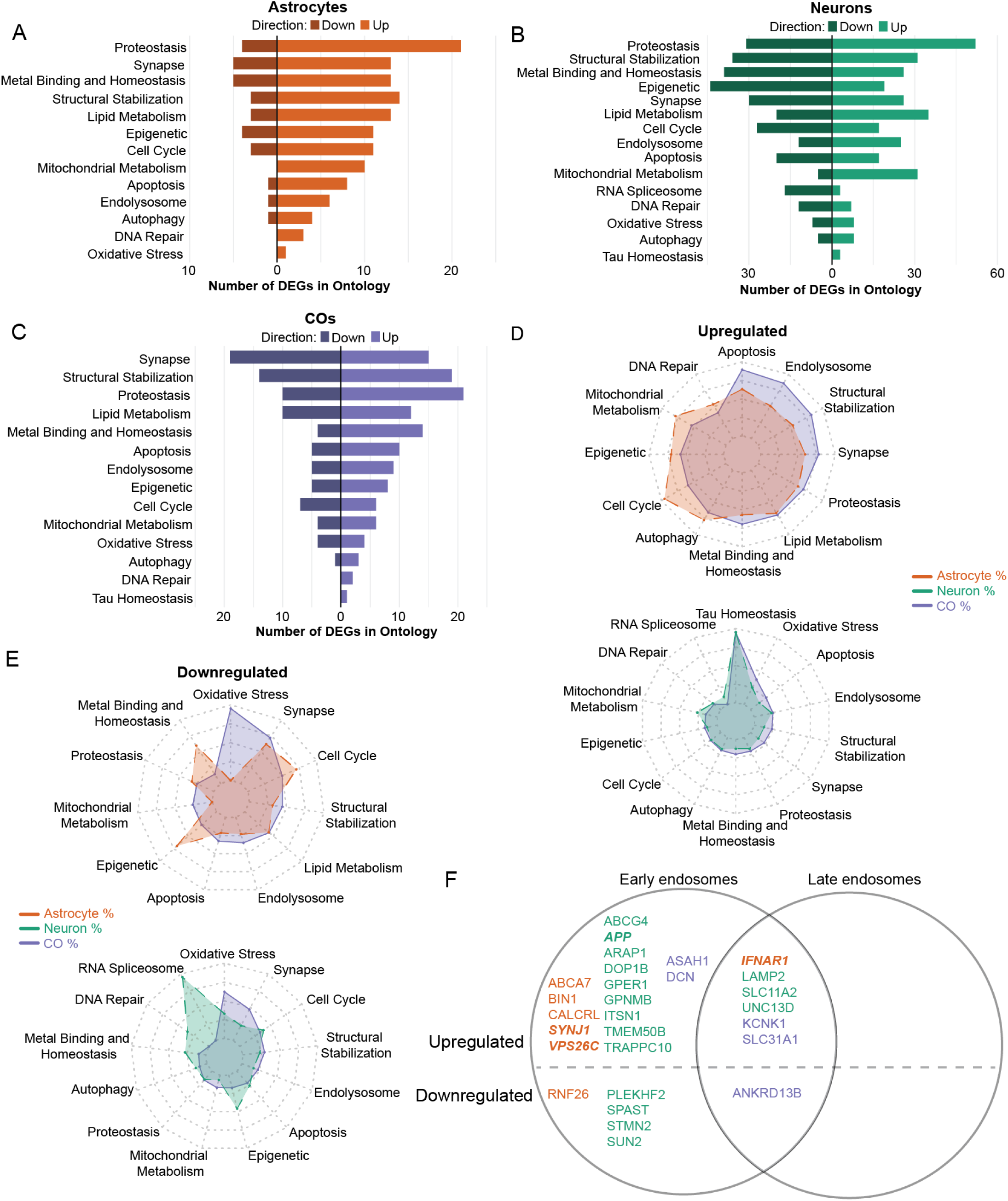
AD pathway-associated gene expression is altered in T21 astrocytes, neurons, and COs. (A) Bar graph showing the number of upregulated (to the right) and downregulated (to the left) significant DEGs between T21 and D21 in hiPSC-derived astrocytes that map to AD-associated ontologies as shown on the y-axis. (B) Bar graph showing the number of upregulated (to the right) and downregulated (to the left) significant DEGs between T21 and D21 in hiPSC-derived forebrain neurons that map to AD-associated ontologies as shown on the y-axis. (C) Bar graph showing the number of upregulated (to the right) and downregulated (to the left) significant DEGs between T21 and D21 in COs map to AD-associated ontologies as shown on the y-axis. (D) Radar plots of upregulated DEGs in COs and hiPSC-derived astrocytes (top) and COs and hiPSC-derived neurons (bottom) sorted into AD-specific ontologies. DEG counts are normalized to the total number of DEGs for the given sample type and to the total number of genes in the list mapping to the relevant AD ontology. In the top plot, each ring represents 0.1475%, with the outermost ring representing 0.59%. In the bottom plot, each ring represents 0.3775%, with the outermost ring representing 1.51%. (E) Radar plots of downregulated DEGs in COs and hiPSC-derived astrocytes (top) and COs and hiPSC-derived neurons (bottom) separately sorted into AD-specific ontologies. DEG counts are normalized to the total number of DEGs for the given sample type and to the total number of genes in the list mapping to the relevant AD ontology. In the top plot, each ring represents 0.13525%, with the outermost ring representing 0.541%. In the bottom plot, each ring represents 0.43725%, with the outermost ring representing 1.749%. (F) Venn diagram showing genes identified in the endolysosomal pathway with roles in early versus late endosome formation, listed by sample type (astrocytes in orange, neurons in green, and COs in purple).

We next evaluated the AD-associated biological domains of downregulated DEGs, which exhibited far less concordance between the three T21 sample types. Synapse and structural stabilization were largely downregulated in all three T21 sample types (Figure 5A–C), and greater proportions of downregulated DEGs were associated with oxidative stress, cell cycle, and lipid metabolism (Figure 5E). T21 neurons exhibited a unique downregulation of genes associated with the RNA spliceosome (Figure 5E). While altered RNA splicing has been reported before in T21 [42], neurons have not been specifically implicated in this process.

Because we identified EV dysregulation as a key pathological feature of T21 COs, we evaluated DEGs associated with the endolysosomal pathway in all three T21 sample types. We discovered that all such DEGs were involved in either early endosomal formation or early/late endosome function, and that Hsa21 DEGs were exclusive to the early endosome (Figure 5F). Notably, *SYNJ1*, *LAMP2*, and *ASAH1* regulate processes ranging from early endosomal maturation to lysosomal trafficking and lipid-dependent EV biogenesis [43–45]. Taken together, these findings suggest early endosomal trafficking dysfunction as a potential link between T21 gene dysregulation and altered EV secretion.

### T21 COs exhibit dysregulation of *APOE*-associated pathways

Finally, we examined DEGs known to interact as either genes or proteins with *APOE* and *APP* as a mechanistic basis for altered apoE and Aβ protein levels in T21 COs. We employed a gene and protein interaction mapping resource [46] and identified 43 genes encoding *APOE*-interacting genes or proteins (8.9% of all DEGs) and 55 genes encoding *APP*-interacting genes or proteins (11.4% of all DEGs) in the three T21 sample types. *APOE*-interacting genes and proteins included upregulated *APP* and *ABCA7* (Figure 6A) and downregulated *SIRT1* (Figure 6B). *APP*-interacting genes and proteins included upregulated *TGFB1* (Figure 6C) and downregulated *SIRT1* and *TTR* (Figure 6D). Most DEGs encoding *APOE*- and *APP*-interacting genes and proteins were unique to T21 neurons, though some, such as *ABCA7*, were unique to T21 astrocytes (Figure 6A–D). We found a high level of confidence (10+ independent confirmations) in the upregulated *ABCA7*, *BIN1*, and *SOD1* genes and in the downregulated *SIRT1* gene as interactors with both *APOE* and *APP* (Figure 6A–D). These genes are likely involved in mechanistic pathways linking *APOE* to DS-AD pathologies in the human brain.

**Figure 6.**
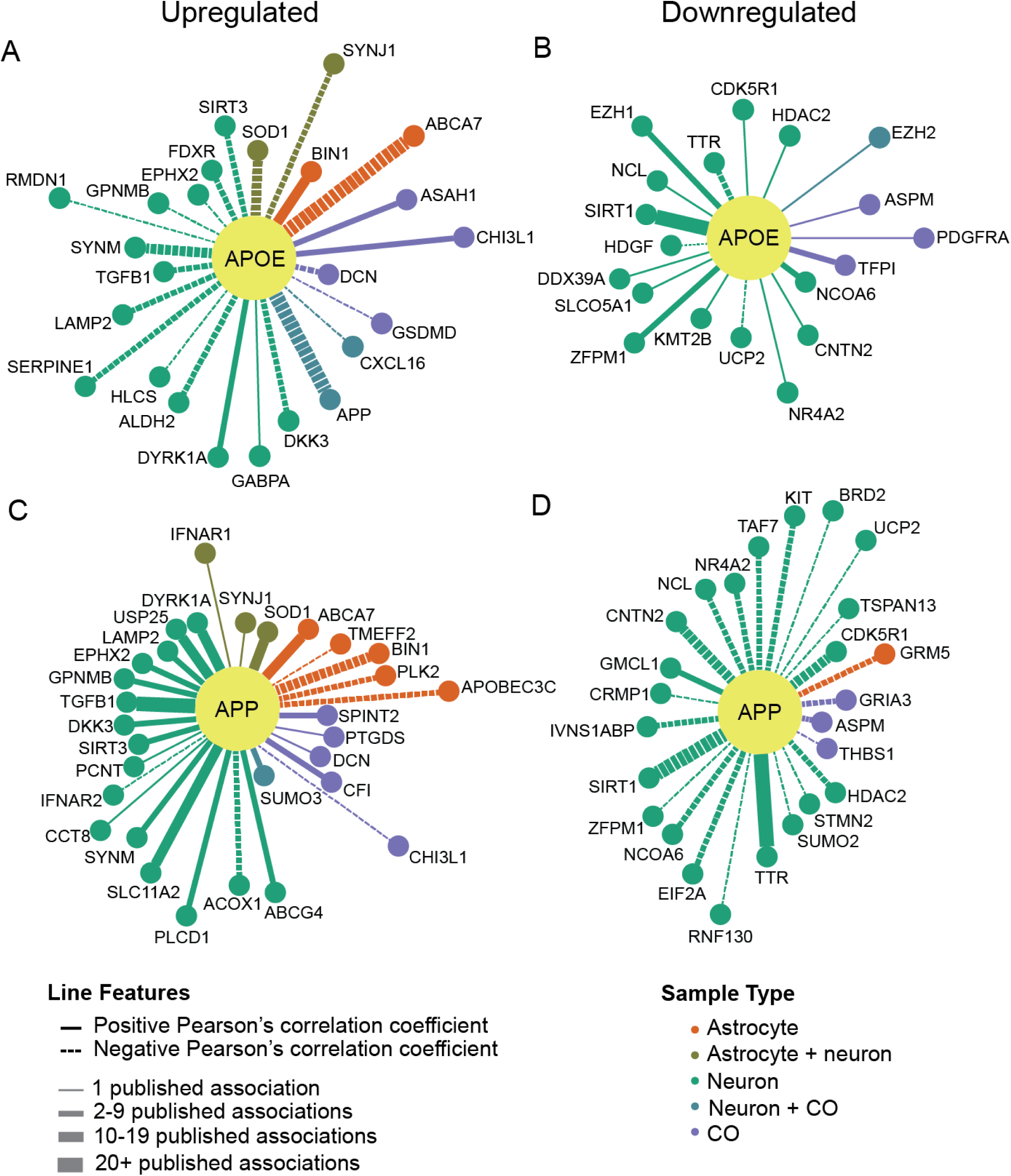
DEGs have documented interactions with APOE and APP. (A) Hub and spoke plot showing binned counts of publications that demonstrate either a gene or protein interaction between *APOE* and upregulated DEGs. Increased line weight indicates a greater number of publications. Distance from the *APOE* node represents Pearson’s correlation coefficient between *APOE* and the DEG as calculated in variance stabilizing transformation (VST)-normalized CO counts. (B) Hub and spoke plot showing binned counts of publications that demonstrate either a gene or protein interaction between *APOE* and downregulated DEGs as discovered using the GePI (Gene and Protein Interactions) web portal [46]. Increased line weight indicates a greater number of publications. Distance from the *APOE* node represents Pearson’s correlation coefficient between *APOE* and the DEG as calculated in VST-normalized CO counts. (C) Hub and spoke plot showing binned counts of publications that demonstrate either a gene or protein interaction between *APP* and upregulated DEGs as discovered using GePI. Increased line weight indicates a greater number of publications. Distance from the *APP* node represents Pearson’s correlation coefficient between *APP* and the DEG as calculated in VST-normalized CO counts. (D) Hub and spoke plot showing binned counts of publications that demonstrate either a gene or protein interaction between *APP* and downregulated DEGs as discovered using GePI. Increased line weight indicates a greater number of publications. Distance from the *APP* node represents Pearson’s correlation coefficient between *APP* and the DEG as calculated in VST-normalized CO counts.

## Discussion

We utilized an *in vitro* model system of the human DS-AD brain to interrogate the individual roles of neurons and astrocytes in the disease process. We identified developmental differences in DS-AD COs compared to euploid COs at the cellular and transcriptional levels that illuminate the role of apoE as a pathological chaperone of Aβ driving AD neuropathology in DS. Given that Aβ polymerization is the key initiating event leading to all other AD-associated pathologies [47, 48], we identified the downstream effects that appear to be mediated, at least in part, by apoE. Our finding that apoE and Aβ protein levels are positively correlated in T21 COs is consistent with studies showing their upregulated levels in the frontal cortex of people with DS [15], and is consistent with the molecular interaction between apoE and Aβ fibrils [49], thereby confirming the overall validity of our model for studying apoE in DS-AD. Finally, our findings highlight the nuanced cell type-specific differences in gene expression that are likely to contribute to established patterns of dysregulation in DS-AD.

We observed an increase in apoE protein levels in T21 COs relative to D21 COs but did not observe a corresponding increase in *APOE* mRNA expression levels by bulk RNA sequencing. This discrepancy could be due to differences in the production, regulation, or utilization of apoE protein in T21 COs, resulting in its accumulation. In the brains of individuals with DS, *APOE* mRNA expression has been found to be increased in several subtypes of astrocytes, endothelial cells, and pericytes [15], which may not be present in our CO model system. The accumulation of apoE protein observed in both T21 COs and in T21 hiPSC-derived astrocytes in lieu of changes in *APOE* mRNA expression indicates that translational control or post-translational mechanisms are likely to contribute, at least in part, to apoE dysregulation in DS-AD.

We also investigated EV release and AD-associated EV cargo from T21 and D21 COs and identified a pattern characterized by increased EV biogenesis signatures together with reduced AD-associated EV cargo levels. Our results showed that EVs released from T21 COs exhibited increased levels of ALIX and altered tetraspanin composition. We further found that cargo levels of Aβ40, Aβ42, tau, and pTauT231 were reduced in T21 EVs compared to D21 EVs. Lower levels of these proteins were also observed in T21 conditioned media. Taken together, our results suggest that T21 is associated with alterations in EV biogenesis and cargo loading pathways.

Previous work has demonstrated that T21 cells exhibit enlarged endosomes and altered endosomal-lysosomal pathway [50]. Increased EV secretion has been reported in T21 brain tissue and cellular models, and this has been proposed to be a compensatory mechanism to impaired endosomal trafficking and cargo accumulation [36, 51]. Our data suggest that T21 COs release greater numbers of EVs, but due to variability in our D21 samples, we were unable to draw a significant conclusion. Additionally, we observed increased ALIX levels in T21 CO-derived EVs. Taken together, these findings support the idea of enhanced activation of EV biogenesis pathways in trisomy 21. We also identified altered tetraspanin distribution in T21 CO-derived EVs, with a reduced proportion of CD63^+^ EVs together with an increased proportion of CD81^+^ EVs, suggesting that T21 influences both EV subtype composition and EV biogenesis pathways. This altered tetraspanin distribution may reflect changes in endosomal maturation, cargo sorting mechanisms, or selective release of distinct EV subpopulations [52, 53]. Because different EV populations can differ substantially in their cargo composition and biological functions, these changes may contribute to altered intercellular signaling and propagation of pathology in DS.

However, despite evidence of increased EV pathway activation, AD-associated proteins were consistently reduced in T21 EVs. These findings suggest that increased EV production is not accompanied by efficient incorporation of AD-associated cargo, which might reflect altered cargo sorting mechanisms or impaired trafficking through endosomal compartments [54].

Notably, reduced levels of soluble Aβ and tau species were also observed in conditioned media from T21 COs. Because the majority of extracellular Aβ and Tau are secreted through non-EV pathways [55], these findings suggest that the trafficking abnormalities observed in T21 are not restricted to EV-mediated export alone. Overall, our results point to a broader dysregulation of intracellular protein trafficking and secretion [56] and are consistent with previous studies suggesting that endosomal abnormalities precede overt extracellular amyloid deposition in DS and AD [50, 57].

In contrast to our findings, Fertan and colleagues reported elevated levels of extracellular soluble Aβ and tau species in conditioned media from T21 organoids using ultrasensitive molecular approaches in which Aβ and tau levels were normalized to metabolic activity based on measures of leftover glucose [58]. In our analyses, we normalized to organoid volume (mm^3^) and found the opposite result. Mitochondrial dysfunction has been extensively reported in DS [59, 60] and is compensated by increased glycolysis to meet energy demands [61]. Therefore, it is possible that, per unit volume, T21 COs consume higher amounts of glucose than D21 COs, which would lead to reduced leftover glucose and therefore inflate normalized Aβ and tau levels in T21 COs. The discrepancy between the results reported herein and those by Fertan et al. likely highlights a confounding artifact introduced by metabolic dysfunction in T21 COs.

Collectively, our findings support a model in which T21 cerebral organoids exhibit increased EV biogenesis but impaired functional secretion of AD-associated proteins across both EV-associated and EV-independent extracellular pathways, suggesting that T21 may initially promote a state of trafficking dysfunction characterized by compensatory activation of EV pathways together with inefficient cargo export. These observations highlight the importance of considering EV biogenesis, cargo loading, and overall extracellular secretion as mechanistically distinct processes and suggest that early alterations in intracellular trafficking may represent a critical initiating event in DS-AD pathogenesis. In support of this, our transcriptomics analysis pointed to altered early endosomal trafficking pathways in T21 COs. Our results indicate that several genes involved in endosomal sorting (*VPS26C*), membrane trafficking and turnover (*SYNJ1*, *ITSN1*, *TRAPPC10*, *TMEM50B* and *ARAP1*), and autophagy (*LAMP2*) are upregulated in T21 COs, pointing to trafficking disruption that occurs during early stages of endosomal sorting.

We additionally detected numerous cell type-specific DEGs that may contribute to apoE-driven DS-AD pathology. First, we observed a ∼1.5-fold upregulation of some, but not all, Hsa21 genes, consistent with prior studies of the DS transcriptome [62, 63]. Second, many DEGs identified across the three sample types (i.e., neurons, astrocytes, and COs) were non-Hsa21 genes, which represent T21-mediated perturbations in gene expression. For example, *H2BC5* (located on chromosome 6) was significantly upregulated in both T21 astrocytes and T21 COs and was previously shown to be involved in nucleosome organization [64] with increased expression in gliomas [65]. The dysregulation of *H2BC5* in glial disease, paired with our identification of its upregulation in T21 models, suggests that it may contribute to a reactive astrocyte-like glial phenotype in DS-AD.

The interactions between *APOE* or *APP* and many of the DEGs identified in this study allowed us to investigate broad patterns of altered pathways that could help explain the changes we observed in apoE and Aβ. Most downregulated DEGs with documented gene or protein interactions with *APP* had inverse associations (negative Pearson’s correlation coefficients) with *APP* gene expression in our dataset, as expected because *APP* was upregulated ∼1.5-fold as an Hsa21 gene. Accordingly, the upregulated DEGs that had gene or protein interactions with *APP* were largely positively correlated. On the other hand, most downregulated DEGs with documented gene or protein interactions with *APOE* had positive associations, and most upregulated DEGs with documented gene or protein interactions with *APOE* had negative associations. Given this, the genes and encoded proteins of DEGs that interacted with both *APP* and *APOE* largely had inverse directionality of association with each of the two genes. Additionally, and notably, most negatively-associated upregulated DEGs that had documented gene or protein interactions with *APP* were astrocyte-specific. As astrocytes are the primary producers of apoE in the brain, these genes may play key roles in the link between *APOE* and amyloid neuropathology in DS-AD.

We identified numerous DEGs that have reported gene or protein interactions with *APOE*, which may provide important insights into mechanisms of DS-AD. *CXCL16* and *EZH2* were differentially expressed in both T21 neurons and T21 COs, despite being non-Hsa21 genes. *CXCL16* (located on human chromosome 17) was previously shown to be protective against neuronal excitotoxicity [66]. *EZH2* (located on human chromosome 7) has been shown to be downregulated at both the gene and protein levels in the AD brain [67], consistent with our findings in DS-AD COs. *EZH2* is involved in polycomb repressive complex 2 (PRC2) activity [68] and determination of cell fate [65, 69], of particular relevance to people with DS whose brains have altered neural cell composition [70]. *ABCA7*, *BIN1*, *SOD1*, and *SIRT1* each had more than 10 reports of interactions with both *APOE* and *APP*. These genes are of particular interest because they may be involved in pathways that underlie the positive correlation in the levels of *APOE* and *APP* proteins observed in T21 COs. *ABCA7* (located on human chromosome 19) is a key lipid transporter that directly influences AD pathology, and genome-wide association studies (GWAS) have determined that loss-of-function mutations in *ABCA7* increase AD risk [71, 72], and *ABCA7* knockout has been shown to increase Aβ levels by approximately two-fold [73]. *BIN1* (located on human chromosome 2) is involved in APP processing [74, 75] and has also been identified previously in GWAS as an AD risk gene [76]. *SOD1* (located on Hsa21) encodes a cytosolic antioxidant enzyme [77] and *SOD1* knockout in an AD mouse model accelerated amyloid plaque deposition [78], illuminating its potential as a therapeutic target for DS-AD. Finally, *SIRT1* (located on human chromosome 10) is a deacetylase that acts on multiple target proteins to slow cellular senescence [79], potentially contributing to the accelerated aging observed in people with DS [80]. This set of genes, along with other DEGs identified in our transcriptomic studies of T21 COs, reflect important pathways implicating apoE in the Aβ-driven pathogenesis of DS-AD, which may be explored mechanistically in the future to identify novel drug targets.

In sum, the present study leverages a powerful *in vitro* model of DS-AD to provide new insights into various mechanistic underpinnings of disease progression. An increased understanding of these mechanisms empowers us to make informed decisions about how to design DS-AD treatments to improve the quality of life for people with DS.

## Methods

### Differentiation of hiPSCs

The generation and validation of hiPSCs were performed as previously described [81] by the Stem Cell Biobank and Disease Modelling Service Core for the Linda Crnic Institute for Down Syndrome at the University of Colorado Anschutz as part of the Human Trisome Project (www.trisome.org, NCT02864108). Astrocytic differentiation was performed according to the STEMdiff™ Astrocyte Differentiation Kit (STEMCELL, Catalog #100-0013), and neuronal differentiation was performed according to the STEMdiff™ Forebrain Neuron Differentiation Kit (STEMCELL, Catalog #08600). Both cell types were grown in 6-well tissue culture-treated plates (Corning, Cat. #353046). COs were generated following established protocols [16] utilizing the STEMdiff™ Cerebral Organoid Kit (STEMCELL, Catalog #08570). After completion of the kit’s protocol (i.e., 40 days *in vitro*), COs were kept in 6-well tissue culture-treated plates (Corning, Cat. #353046) in 3 mL of maintenance media from the Maturation Kit (STEMCELL, Catalog #08571) with media changes every three days until 120 DIV. We maintained approximately 10 COs in each well of a 6-well plate.

### Astrocyte and neuron fixation and immunofluorescence microscopy

Monolayer cell cultures were prepared for immunofluorescence staining in 96-well polymer coverslip plates (Ibidi, Catalog #89626). Cells were washed once in 1x phosphate-buffered saline (PBS) and then fixed in 150 µL of 4% paraformaldehyde (PFA) (Electron Microscopy Sciences 15700-1L; diluted in Milli-Q water) for 15 minutes at room temperature. Cells were then washed three times with 1X PBS for five minutes each. To permeabilize and block, 150 µL of filtered (EMD Millipore, Cat. #SCGP00525) 5% BSA (EMD Millipore, Cat. #2930-100GM) with 0.1% triton-X (made in 1X PBS) was added to each well and incubated for one hour at room temperature. Monolayer astrocytes were stained with antibodies against apoE (Abcam, Cat. #ab183597, 1:500) and GFAP (Abcam, Cat. #ab4674, 1:1000) diluted in 5% BSA. Monolayer neurons were stained with antibodies against MAP2 (antibodiesinc, Cat. #1100-MAP2, 1:1,000) and NeuN (Abcam, Cat. #ab104224, 1:100) diluted in 5% BSA. For each cell type, 150 µL of diluted antibody was added per well and incubated at 4°C overnight.

The following day, all wells were washed four times with 1X PBS five minutes each. The slides were then incubated in 200 µL of fluorescent secondary antibodies for wavelengths 488 nm (Fisher Scientific, Cat. #PIA32723, 1:250), 555 nm (Fisher Scientific, Cat. #PIA32732, 1:250), and 647 nm (Fisher Scientific, Cat. #A21449, 1:250) were incubated at room temperature for one hour. Following this, all wells were washed twice with 1X PBS for five minutes each. Nuclear stain (Fisher Scientific, Cat. #PI62249, 1:12,500) diluted in 1X PBS was then added to each well and incubated for 10 minutes at room temperature. Wells were then again washed twice with 1X PBS for five minutes each. After washing, 200 µL of 1X PBS was added to each well, and plates were taken for imaging.

### CO fixation and cryosectioning

COs harvested for immunofluorescence microscopy were removed from their 6-well plate and immediately placed individually into 500 µL of 1X PBS in a 48-well plate (Corning, Cat. #351178). All 1X PBS was then removed, and 500 µL of 4% PFA (diluted in Milli-Q water) was added to each well for fixation. Following a 1-hour fixation at RT, all 4% PFA was removed from each well, and 500 µL of 1X PBS was added for five minutes to wash. COs were then taken through a stepwise sucrose gradient (6.25% for 45 min, 12.5% for 3-4 hours, and 25% overnight). For each step, 500 µL of the sucrose solution (D-sucrose, Fisher Scientific, Cat. #BP220-212, diluted in 1X PBS) was incubated at room temperature for the indicated time or until all COs had sunk to the bottoms of their wells. The following day, each CO was removed from sucrose solution and transferred to a disposable base mold (Fisher Scientific, Cat. #22-363-553). Excess sucrose solution was removed from the base mold, and the mold was filled with Tissue-Tek^®^ O.C.T. Compound (Sakura, Cat. #4583) and left for one hour at room temperature. Following this, each entire base mold containing O.C.T. and COs was flash-frozen by placing it in liquid nitrogen-chilled isopentane (GFS Chemicals, Cat. #46085) until the O.C.T. turned completely solid and white in color. Frozen molds were stored at -80°C until they were sliced into 20 µm-thick sections at -20°C using a cryostat and mounted onto positively-charged microscopy slides (Electron Microscopy Sciences, Cat. #71869-11). Slides with CO sections were stored at -80°C until immunofluorescent staining.

### CO immunofluorescence staining and microscopy

In preparation for immunofluorescent staining, slides were removed from their -80°C storage and placed at room temperature to defrost. Slides were then submerged in 1X PBS for five minutes to wash then quickly dipped into deionized water before being transferred to a Sequenza™ Immunostaining Center Slide Rack (Epredia, Cat. #73310017). Slides were permeabilized for 10 min using 0.1% Triton X-100 (diluted in 1X PBS) at room temperature, then washed with 1X PBS for five minutes at room temperature. Then, 1 mL of filtered (EMD Millipore, Cat. #SCGP00525) 5% BSA (EMD Millipore, Cat. #2930-100GM) (made in 1X PBS) was added to fill each slide rack chamber and incubated for one hour at room temperature. Following this, antibodies against MAP2 (antibodiesinc, Cat. #1100-MAP2, 1:1,000), apoE (Abcam, Catalog #ab183597, 1:500), and Aβ (BioLegend, Cat. #803001, 1:250) were diluted in filtered 5% BSA and 200 µL was added to each slide rack chamber to incubate at 4°C overnight.

Slides were washed with 1X PBS four times (5 minutes each). Thereafter, the slides were incubated in 200 µL of fluorescent secondary antibodies for the wavelengths 488 nm (Fisher Scientific, Cat. #PIA32723, 1:250), 555 nm (Fisher Scientific, Cat. #PIA32732, 1:250), and 647 nm (Fisher Scientific, Cat. #A21449, 1:250) diluted in 5% BSA for one hour at room temperature. Slides were then washed twice in 1X PBS. Then, 200 µL of nuclear stain (Fisher Scientific, Cat. #PI62249, 1:12,500) diluted in 1X PBS was added to each chamber and incubated for 10 minutes at room temperature. Wells were again washed twice with 1X PBS for five minutes each. Slides were then removed from the slide rack chamber and quick-dried. Immediately after, 100 µL of hard-set mounting media (Thermo Scientific, Cat. #P36934) was added to the top of each slide using a wide-bore 200 µL pipette tip (Fisher Scientific, Cat. #14-222-730), and a glass cover slip (Corning, Cat. #2975-225) was applied to cover the mounting media and the tissue. Slides dried for 48 hours at room temperature and then were taken for imaging.

### Protein extraction and lysate preparation

COs harvested for immunoblotting were removed from their 6-well plate and immediately placed individually in 500 µL of 1X PBS in a 48-well plate (Corning, Cat. #351178) to wash. Each CO was then transferred to a 1.7 mL microcentrifuge tube (Fisher Scientific, Cat. #MCT-175-L-C) on wet ice. Excess 1X PBS from the transfer step was removed, and 100 µL of RIPA buffer (Fisher Scientific, Cat. #PI89901) with protease inhibitors (Thermo Scientific, Cat. #78442) was added to each microcentrifuge tube. COs were then sonicated for 10 seconds each at 100% amplitude and incubated on ice for 30 minutes. Samples were spun at 20,000 x g for 10 minutes at 4°C to separate soluble from insoluble lysates. Soluble lysates were transferred to individual wells of a 96-well polypropylene plate (USA Scientific, Cat. #1833-9610). Then, 8 µL of each sample was transferred to a new polypropylene plate and diluted five-fold in 32 µL of 1X PBS. To determine the concentration of each sample, a BCA assay was performed (Thermo Scientific, Cat. #23227) according to kit instructions and using 10 µL of each diluted sample in a 96-well polystyrene plate (Thermo Scientific, Cat. #442404). All samples were then normalized by concentration by diluting samples with higher concentrations in RIPA buffer with protease inhibitors. Normalized samples were stored as 15 µL aliquots in polypropylene plates sealed with aluminum seal tape (Thermo Scientific, Cat. #232698) at -80°C.

### Immunoblotting

BCA-normalized CO protein lysates were thawed on ice until fully liquid. In a PCR plate (BioRad, Cat. #HSP9601B), 15 µL protein lysate (> 10 µg total protein), 5.75 µL water (Thermo Scientific, Cat. #R0581), 6.25 µL LDS sample buffer (Invitrogen, Cat. #NP0007), and 3 µL reducing reagent (Invitrogen, Cat. #NP0009) were combined and incubated at 95°C for 10 minutes. While the reducing reaction was occurring, a precast protein gel (Thermo Scientific, Cat. #WG1403BX10) was rinsed with deionized water, its comb and tape were removed, and it placed in a Criterion™ electrophoresis cell (BioRad, Cat. #1656001). Running buffer concentrate (Invitrogen, Cat. #NP0001) was diluted in Milli-Q water and added to fill the electrophoresis cell. To ensure the samples stayed submerged in running buffer, a Midi Gel Adapter (Thermo Scientific, Cat. #WA0999) was added to the cassette holding the precast gel. To this adapter was added 50 mL of running buffer with 25 µL of antioxidant (Invitrogen, Cat. #NP0005). Each well of the precast gel was rinsed with the antioxidant-containing running buffer prior to sample loading. 10 µL of protein ladder (Thermo Scientific, Cat. #26616) or of sample was added per well, and the gel was run at 200 V for 45 minutes (or until the gel front had migrated to the bottom of the gel). To transfer the gel to a membrane, a prepared PVDF transfer stack (Thermo Scientific, Cat. #IB24002) was placed in an iBlot™ 2 Gel Transfer Device (Thermo Scientific, Cat. #IB21001). Approximately 2 mL of Milli-Q water were added on top of the membrane, and the completed gel was placed on this liquid. The PVDF stack was reassembled, and air bubbles were removed with a roller (Invitrogen, Cat. #LC2100). The iBlot™ machine was then run at 20 V for 7 minutes to complete the transfer. The membrane was then removed from the PVDF stack and placed into 15 mL of 3% BSA in 1X tris-buffered saline (TBS) (Thermo Scientific, Cat. #28358, diluted to 1X in Milli-Q water) with Tween^®^ 20 (Fisher Scientific, Cat. #BP337500) (TBS-T) added to a final working concentration of 0.01%. The membrane was left to block in this solution at room temperature on a shaker on low setting for one hour. Following this, the blot was divided and separately probed with primary antibodies against apoE (Abcam, Cat. #ab183597; 1:1,000) or Aβ (BioLegend, Cat. #800701, 1:500). These antibodies were added directly to the blocking solution, and the solution with the membrane was placed on a low-setting shaker at 4°C to incubate overnight.

The following morning, all the primary antibody solution was emptied from around the membrane, and the membranes were washed four times with 1X TBS-T for five minutes each on the shaker at room temperature. Then, secondary antibody dilutions were prepared in 15 mL of 3% BSA in TBS-T. Anti-mouse IgG (Jackson ImmunoResearch, Cat. #115-035-003, 1:5,000) was added to the blot probed for Aβ, and anti-rabbit IgG (Jackson ImmunoResearch, Cat. #111-035-144, 1:5,000) was added to the blot probed for apoE. Secondary antibody solution was incubated with the blots on a shaker at room temperature for one hour. Each blot was then washed an additional four times with TBS-T for five minutes each. Chemiluminescence reagents (Thermo Scientific, Cat. #34577) were then mixed at a 1:1 ratio, and each blot was incubated in this solution for 90 seconds at room temperature. Immediately following, blots were wrapped in plastic film and taken for imaging on an iBright™ Imaging System.

To normalize the protein levels from sample to sample, each blot was stripped and re-probed for β-actin. Stripping solution (EMD Millipore, Cat. #2504) was diluted to a 1X working concentration in Milli-Q water. Blots were placed in the diluted stripping solution and incubated on a shaker for 10 minutes at room temperature. After this, each blot was washed once with 1X TBS-T. Blocking was then performed again using 15 mL of 3% BSA in TBS-T for one hour at room temperature. Antibody against β-actin (Santa Cruz Biotechnology, Cat. #sc-47778, 1:2,000) was then added directly to the blocking solution of each blot, and the incubation was moved to 4°C to carry out on a low-setting shaker overnight.

The following day, the wash steps were repeated, and anti-mouse IgG (Jackson ImmunoResearch, Cat. #115-035-003, 1:5,000) secondary antibody was diluted in 3% BSA in TBS-T and incubated for one hour at room temperature. The remaining wash and chemiluminescence steps were repeated prior to imaging.

### EV enrichment from CO media

Conditioned media from two biological replicates of either D21 or T21 isogenic hiPSCs line-derived COs were collected at two time points (60 DIV and 120 DIV post-differentiation). Three technical replicates were processed for each of the four 60 DIV CO lines, providing a total of 12 samples (n=6 T21 and n=6 D21). For the older (120 DIV) COs, 3-4 replicates were used per CO line, yielding 7 and 9 technical replicates for the T21 and D21 COs, respectively. Cell culture media was first centrifuged at 3,000 x g for 20 minutes at 4°C to remove cell debris. The supernatant was filtered through a 0.22-μm pore PES membrane (SCGP00525; Millipore). Five mL of cleared conditioned media was concentrated on a 10 kDa MWCO filter device (UFC8010; Millipore) preconditioned with sterile-filtered Dulbecco’s PBS (DPBS). Media was washed once by adding DPBS to the filter device, followed by centrifugation (3,000 x g, 15 min, 4°C). The resulting sample was transferred into a clean microtube, and the volume was brought to 1 mL by adding DPBS. EVs were precipitated overnight at 4°C with 200 μL of ExoQuick-TC (EXOTC50A-1; SBI) and then centrifuged (1,500 x g, 30 min, 4°C). The pelleted EVs were resuspended in 300 μL DPBS. Intact EVs were aliquoted and stored at -80°C until use.

### EV characterization

Nanoparticle tracking analysis was performed using a ZetaView PMX 120 V4.1 (Particle Metrix, Germany). CO-derived EVs (10-20 μL) were fluorescently labelled with αCD9 αCD63 αCD81 antibody solution (Particle Matrix) at a 1:10 final dilution overnight at 4°C. Sample volumes were brought up to 1 mL with DPBS and directly injected for size and concentration assessment in fluorescent mode (488 nm). ZetaView parameters were set up for gain/shutter 80/100 for fluorescence (FL500) readings. CO-derived EV morphology was assessed by Transmission Electron Microscopy (TEM). Briefly, 10 µl of EV suspension (in DPBS) were loaded onto a discharged grid and incubated for 5 min at room temperature. The excess liquid was then blotted with filter paper and the grid immediately rinsed with 10 µL of deionized water. Thereafter, 10 µL of filtered 2% aqueous uranyl acetate was added for negative staining. After full removal of the liquid from the grid, samples were air-dried for approximately 3 minutes. Data were collected using the JEOL JEM-120i, 120 kV TEM equipped with an AMT NanoSprint 15Mk-II sensor.

The single-particle interferometric reflectance imaging sensor-based (SP-IRIS) analyzer (ExoView R200, NanoView Biosciences) was used to examine the tetraspanin distribution of CO-derived EVs with the Human Tetraspanin Plasma assay kit (Unchained Labs, Cat. #251-1045) according to the manufacturer’s instructions. This assay captures EVs expressing the tetraspanins CD63, CD81, CD9, with mouse IgG as negative control and CD41a as indicator of platelet-derived EVs [82]. The numbers of sized particles (50-200 nm) were analyzed with the ExoView Analysis software.

CO-derived EVs (5-10 μL) enriched from conditioned media were mixed with 2x Laemmli Sample Buffer containing 5% 2-β-mercaptoethanol and boiled for 5 minutes at 95°C. Samples were loaded on 4–15% Mini-PROTEAN^®^ TGX™ Precast Protein Gels (Bio-Rad, Cat. #4561083) and transferred on 0.2 µm PVDF membranes (Bio-Rad, Cat. #1704156) using a Trans-Blot Turbo Transfer System. Membranes were blocked in blocking buffer (5% low fat milk reconstituted in 0.1% TBS-T for 2 hours and probed overnight at 4°C with primary antibodies against ALIX (System Biosciences, Cat. # EXOAB-ALIX-1, 1:1000), CD81 (Unchained Labs, clone JS-81, 1:200), CD9 (Unchained Labs, clone HI9a, 1:200) and CD63 (Unchained Labs, clone H5C6, 1:200) diluted in blocking buffer. Membranes were washed 3 times in 0.05% TBST, incubated for 1 hour with HRP-conjugated secondary antibodies (Jackson Labs, Cat. #115-035-146 and 111-035-144), developed using ECL substrate and imaged on an iBright Imaging Systems.

### Biomarker analyses

Intact EVs were lysed on ice for 5 minutes with 1.8 volume of M-PER lysis buffer supplemented with 25 μL of 3% BSA and 1X protease and phosphatase inhibitor cocktail (Thermo Scientific, Cat. #78443) and stored at -80°C. Before analysis, the lysed EVs were thawed for 3 min in a 37°C water bath then returned to -80°C for 2 hours and thawed one more time before being centrifuged at 10,000 x g for 5 minutes and aliquoted into small tubes.

The levels of ALIX were measured in all EV preps using a colorimetric ELISA kit (Cusabio, Cat. #EL017673HU) according to manufacturer’s instructions. Levels of Tau (N-terminal to mid-domain (R1) Tau), Aβ40, and Aβ42 were measured in both lysed EV preps (diluted 4-fold) and in media (diluted 200-fold) using the human Neurology 3-plex A assay (Quanterix, MA). Levels of pTau T231 were measured with the Advantage PLUS pTauT231 assay kit (Quanterix, MA). Assays were run according to the manufacturer’s instructions.

## Statistical analyses

Morphological, protein, and EV data were analyzed in GraphPad Prism (version 10). Unpaired two-tailed Student’s t-tests were used to compare D21 and T21 groups, except for CO volume (Figure 1K), which was compared using a 2-way ANOVA. Correlation values were determined by computing Pearson’s correlation coefficient. Data are represented as mean ± standard deviation unless stated otherwise.

## RNA sequencing and analysis

RNA was extracted from astrocytes, neurons, and COs according to the instructions in the AllPrep DNA/RNA/Protein Kit (Qiagen, Catalog #80004). Prior to extraction, all samples had been collected and washed in 1X PBS before being frozen at -80°C in RNAlater (Sigma-Aldrich, Catalog #R0901-100ML). Additionally, COs were homogenized using a QIAshredder (Qiagen, Catalog #79654). During this step, three to five COs for each cell line were pooled to ensure adequate material for sequencing. Samples with RIN ≥ 5.0 and ≥ 1 ng/µL of RNA were sequenced. Samples were sequenced using the Illumina NovaSEQ X as paired-end 2 x 150 bp reads with a sequencing depth of 40 million read pairs per sample.

Raw sequencing counts were subject to quality control using FastQC. No poor-quality reads were identified. Adapter sequences were trimmed from the data using cutadapt/4.2. We then re-performed quality control with FastQC, and no poor-quality reads were identified. Samples were then aligned to the GRCh38 human genome with v46 primary annotation from GENCODE. Alignment was performed using star/2.7.10b. Picard (v2.27.5) was used to collect RNA-seq and alignment metrics for quality control, including the determination of library strandedness. Based on strandedness metrics from Picard, the data were determined to be in fr-secondstrand (F1R2) orientation. Accordingly, strand-specific raw gene counts were extracted from the third column of STAR’s reads per gene output and used for downstream normalization and analysis. RNA sequencing metrics were generated using CollectRnaSeqMetrics from PicardTools. Lowly expressed genes were filtered out by removing those with fewer than 0.25 counts per million in more than 50% of the samples.

To avoid artificial inflation of power, technical replicate samples generated from the same starting cell line were collapsed into one sample by averaging the count tables. Additionally, RUVr was performed on all samples. The surrogate variables W_2 and W_3 were used in neuron and astrocyte analyses, and W_1 was utilized in cerebral organoid analyses. Differential expression analysis was performed using DESeq2.

AD ontologies were applied using previously established gene lists [41]. Associations between DEGs and *APOE* or *APP* were determined as the count of publications linking each DEG to either *APOE* or *APP* as identified using GePI [46].

## Acknowledgements

Research reported in this publication was supported by the National Institute on Aging of the National Institutes of Health (NIH) under award numbers F31AG084295, RF1AG070153, R01AG078965, RF1AG078965, and U24AG092191 and by the NIH Office of the Director under award number R24OD035579. Additional funding support was provided by the Linda Crnic Institute for Down Syndrome and the Global Down Syndrome Foundation. Projects were completed in collaboration with the Drug Discovery and Development Shared Resource at the University of Colorado Anschutz Medical Campus, RRID:SCR_021986, P30-CA046934, ALSAM Foundation, CU Anschutz Center for Drug Discovery. This study was supported by the Electron Microscopy Core Facility at CU Anschutz. Data were collected using the JEOL JEM-120i, 120 kV TEM, supported by NIH grant 1S10OD036258-01.

## Notes

### Competing Interest Statement

The authors have declared no competing interest.

