## Supplemental Materials for "Trisomy 21 cerebral organoids exhibit Alzheimer’s disease amyloid and apolipoprotein E co-pathologies"

D21-1

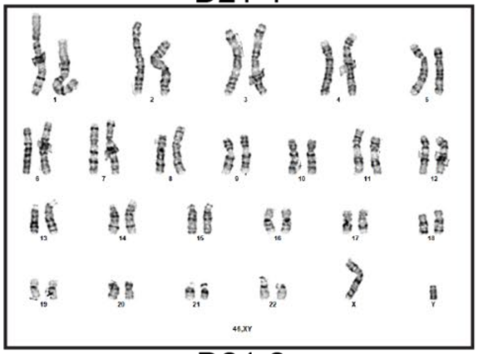


D21-6

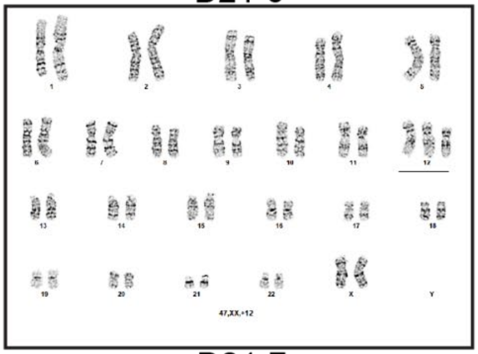


T21-3

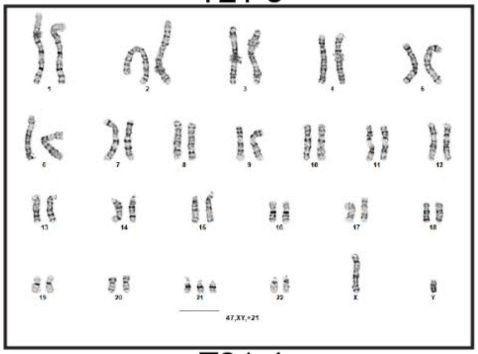


D21-2

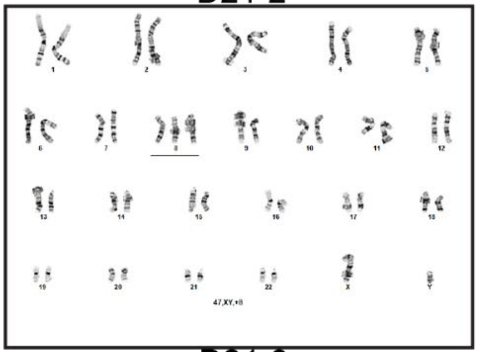


D21-7

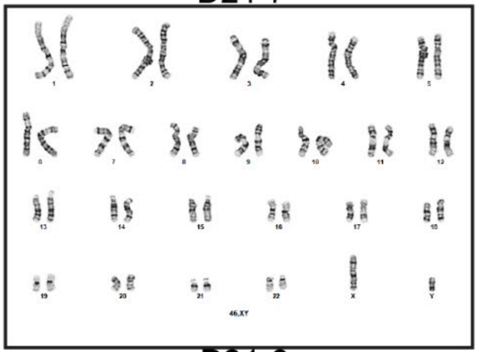


T21-4

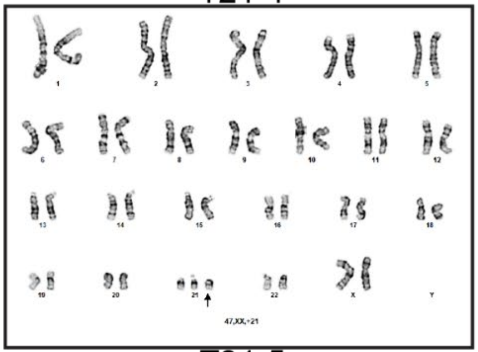


D21-3

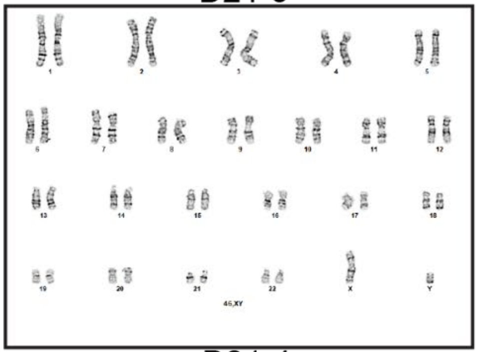


D21-8

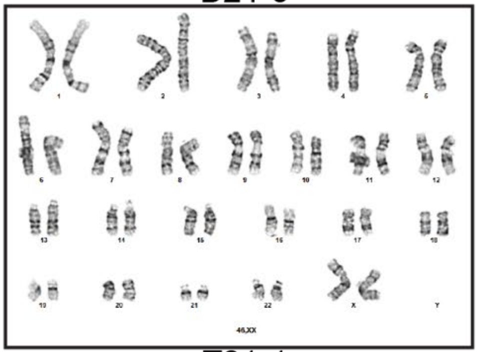


T21-5

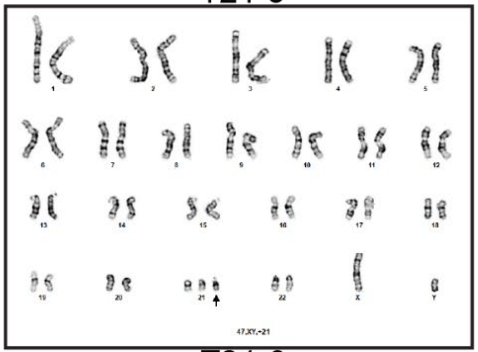


D21-4

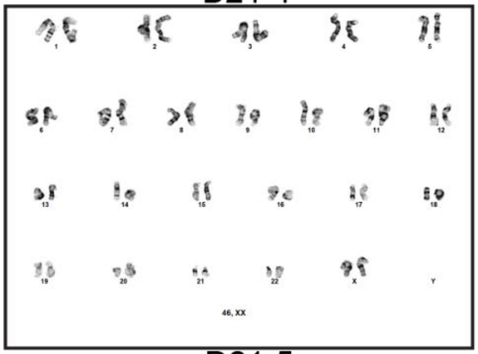


T21-1

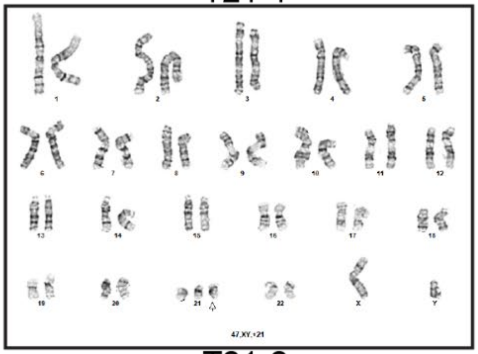


T21-6

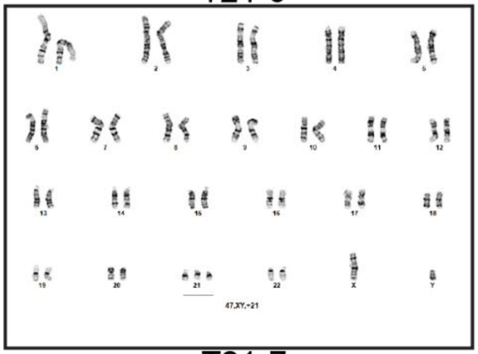


D21-5

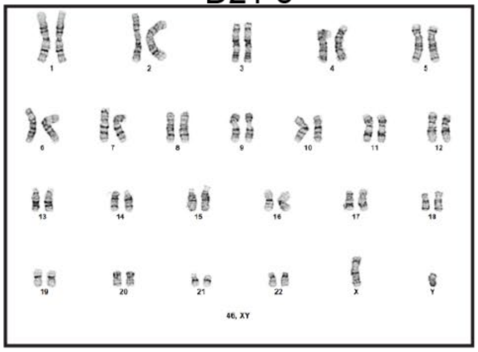


T21-2

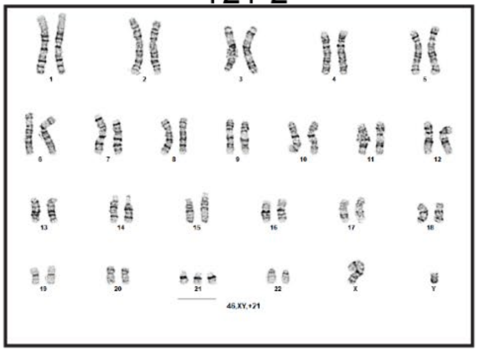


T21-7

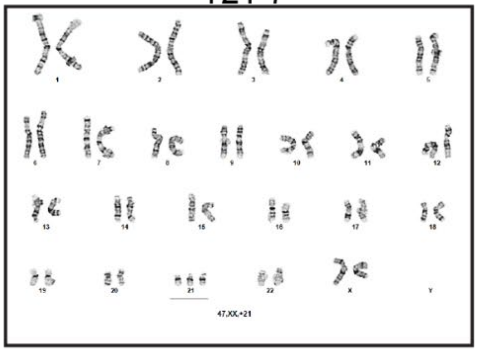

**Supplemental Figure 1. Karyotyping of 15 hiPSC lines.** Karyotypes of each of the 15 hiPSC lines used for all experiments as generated by chromosome harvest and G-banding karyotype analysis. All karyotyping was performed by the Department of Pathology/Colorado Genetics Laboratory within the University of Colorado Cancer Center at the University of Colorado Anschutz Medical Campus.

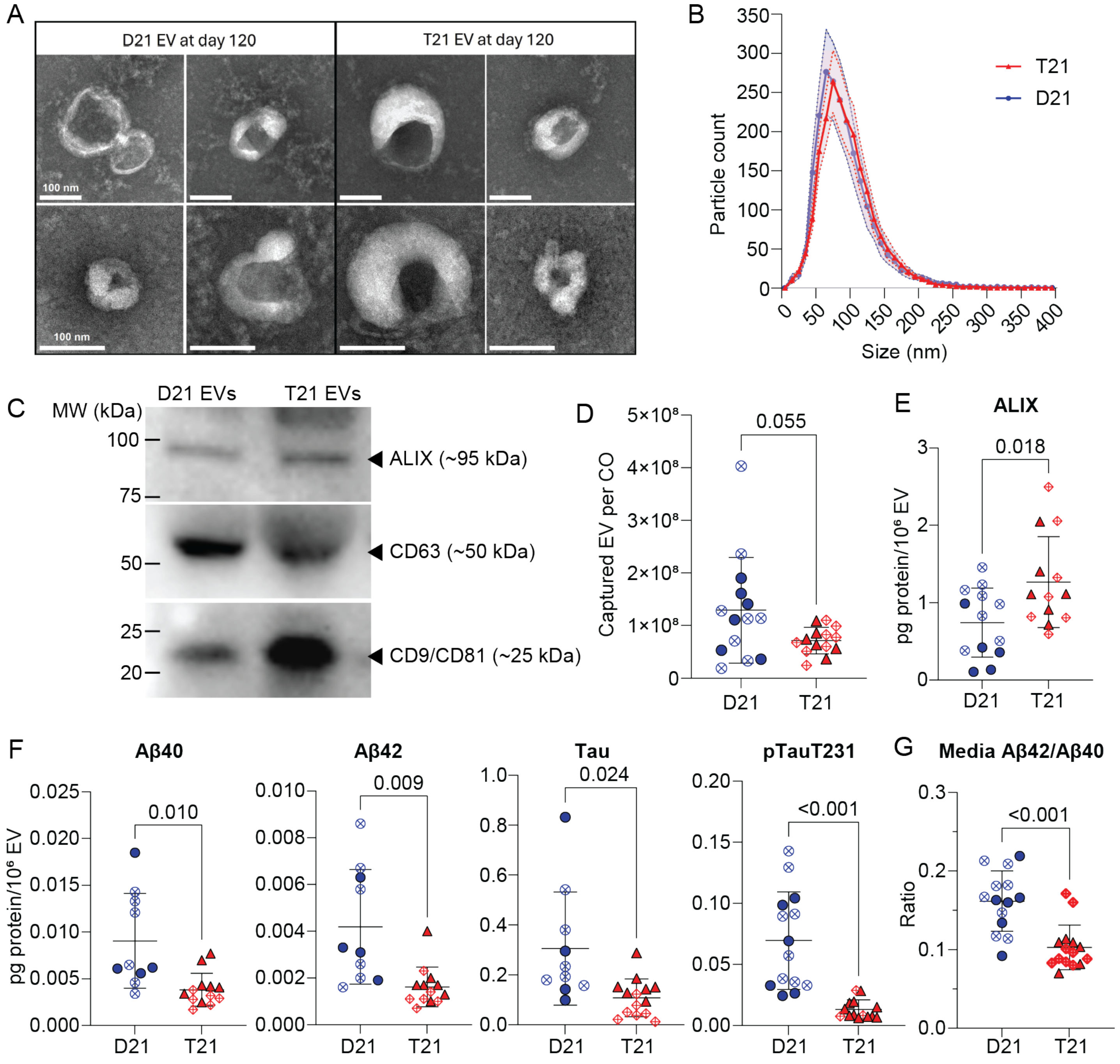


**Supplemental Figure 2. Additional characterization of CO-derived EVs and EV cargo.** (A) Transmission Electron Microscopy (TEM) images of CO-derived EV preps. Samples were imaged on a JEOL JEM-120i, 120 kV TEM equipped with an AMT NanoSprint 15Mk-II sensor. Scale bars represent 100 nm. (B) Nanoparticle tracking analysis (NTA, ZetaView) of fluorescently labelled EVs from D21 and T21 cerebral organoids. Solid lines represent the average particle count from n=6 replicates. Shaded areas represent the standard errors of the mean for each group. (C) Qualitative immunoblot analysis of EV markers ALIX, CD63, and CD9/CD81 in EV preps from conditioned media of 100 DIV euploid (D21) and trisomic (T21) cerebral organoids. Equal amounts of protein were separated on 4-20% SDS-PAGE gel and immunoblotted with anti-human antibodies. (D) Total number of tetraspanin-positive EVs released per CO. EVs were isolated from 5 mL of conditioned media from euploid (D21, n=14) and trisomic T21 (n=14) COs. EVs were incubated overnight on chips coated with anti-CD63, anti-CD81 and anti-CD9 capture antibodies from the Tetraspanin Plasma kit (NanoView). A mouse IgG chip was included as a negative control (not shown). After washing and addition of detection antibodies (anti-CD81-CF555, anti-CD9-CF488, and anti-CD63-CF647) according to the manufacturer's instructions, the chips were scanned on the ExoView R200 instrument. Data are represented as mean of total sized (50-200 nm) captured EVs ± standard deviation. (E) ALIX protein concentration (pg) expressed by $10^6$ EVs. (F) Aβ40, Aβ42, total tau, and pTau231 protein concentrations (pg) expressed by $10^6$ EVs. (G) Ratio of media concentrations of Aβ42 over Aβ40. Data are represented as mean ± standard deviation.

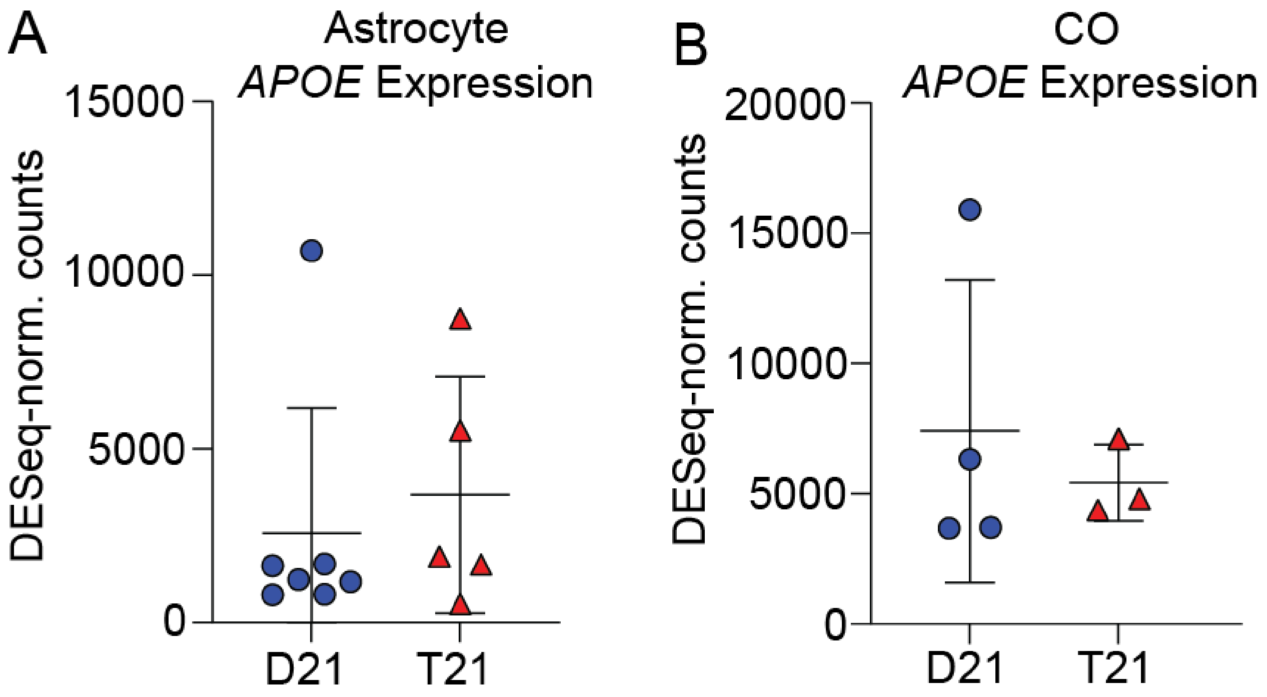


**Supplemental Figure 3. Similar expression levels of *APOE* mRNA in T21 and D21 hiPSC-derived astrocyte and CO models.** (A) DESeq-normalized counts of *APOE* expression as generated by differential expression analysis in hiPSC-derived astrocytes. (B) DESeq-normalized counts of *APOE* expression as generated by differential expression analysis in COs.
